# Mitochondrial Haplogroup Association with Fasting Glucose Response in African Americans Treated with a Thiazide Diuretic

**DOI:** 10.1101/2021.10.18.464878

**Authors:** Bre A. Minniefield, Nicole D. Armstrong, Vinodh Srinivasasainagendra, Hemant K. Tiwari, Scott W. Ballinger, Zechen Chong, Stella Aslibekyan, Donna K. Arnett, Marguerite R. Irvin

## Abstract

Hypertensive African Americans have ~50% response rate to thiazide diuretic treatment. This contributes to a high prevalence of uncontrolled high blood pressure. Here, we examine the role the mitochondrial genome has on thiazide diuretic treatment response in hypertensive African Americans enrolled in a clinical trial. Participants from the Antihypertensive and Lipid Lowering Treatment to Prevent Heart Attack Trial (ALLHAT, n= 4279) were genotyped using the Illumina Infinium Multi-Ethnic Beadchip. Haplotype groups were called using HaploGrep. We used linear regression analysis to examine the association between mitochondrial haplogroups (L, M, and N) and change in blood pressure and change in fasting glucose over 6 months and two years, respectively. The analysis revealed a null association between mitochondrial haplogroups M and N vs. L for each of the outcomes. In subgroup analysis, the L subclades L1, L2, and L3/L4 (vs. L0) were each inversely associated with fasting glucose response (p < 0.05). This discovery analysis suggests the mitochondrial genome has a small effect on fasting glucose but not blood pressure response to thiazide diuretic treatment in African Americans.

## Introduction

Hypertension (HTN) is a chronic condition leading to various deleterious health concerns including cardiovascular disease (CVD).^1^ Reports state African Americans (AAs) have the highest prevalence of HTN (54%) compared to all other American populations (European Americans (46%), Asian Americans (39%), Hispanic Americans (36%), and African-born immigrants (<40%)).^2^ They also have the lowest prevalence of controlled HTN among hypertensive patients receiving medication.^2^

These disparities may in part be due to differences in response to first-line antihypertensive (AHT) medications. It has been established that AAs respond better to diuretics and calcium channel blockers as opposed to other classes of AHT pharmaceuticals.^3–4^ Still, a wide distribution of blood pressure (BP) change on diuretic treatment has been observed, demonstrating the need to understand factors associated with treatment response.^5^ In particular, genetic factors associated with BP response to thiazide diuretics in AA populations are understudied. Moreover, thiazide diuretics are significantly associated with metabolic side effects, like impaired fasting glucose (FG).^5^ Pharmacogenomic studies can offer a solution to some of these problems by identifying genetic contributors to both treatment response and adverse effects. Remarkably, genetic risk factors for BP response to thiazide diuretics are few overall, especially in the AA population.^6^

Mitochondria are cellular energy production organelles so important to cell function that they have their own independent genome. Importantly, mtDNA variation can contribute to an imbalance in cellular energy homeostasis and increase vulnerability to oxidative stress, which has been linked to cardiometabolic diseases like HTN.^7–10^ The mitochondrial genome shows noteworthy diversity, due to a higher mutation rate than the nuclear genome, where most variants induce small changes in mitochondrial function that may contribute to disease.^11,12^ Relevant to our study, previous reports have found several mitochondrial genetic variants associated with systolic BP, FG, and other cardiometabolic traits.^12–14^

Another unique aspect of the mitochondrial genome is that, due to maternal inheritance and lack of recombination events, it displays distinct regional variation (potentially due to natural selection pressures) defining distinct lineages associated with the significant global ethnic groups.^15–19^ Particularly, macrohaplogroup L is specific to sub-Saharan Africa, with the most ancient mtDNA haplogroups being L0 and L1 which form the basal lineages of the human mtDNA tree, followed by L2 and L3.^19^ Previous studies suggest that specific mtDNA haplogroups, especially those specific to sub-Saharan Africa, may be associated with diseases of metabolism in modern times such as CVD, type 2 diabetes (T2D), and HTN.^7, 19, 20^ Further, cybrid studies comparing European-origin haplogroups (H and J) and African-origin haplogroups (L) have shown differences in cell growth, oxygen consumption rates, expression of non-energy-related genes, responses to stressors, and rates of glycolysis.^21, 22^

While mitochondrial dysfunction due to mtDNA variation has been consistently associated with cardiometabolic disease, few studies have examined how variants are associated with response to common treatments for these diseases (including AHTs) despite mounting evidence that the mitochondrial genome can play a role in drug response.^23^ Additionally, most drugs used in cardiology have some degree of mitochondrial toxicity which could be exacerbated by genomic variations.^24^ For example, the use of thiazide diuretics was found to decrease mitochondrial function by inducing apoptosis.^20, 25^ Also, a recent study observed links between nuclear variants with mitochondrial function and drug response to other AHT classes.^26^ Given established racial and inter-individual differences in response to thiazide diuretics, we investigated whether mtDNA variants and haplogroups give additional insight into BP and FG response to thiazide diuretics treatment in AAs.

## Materials and Methods

### Sample Population

All data used in this study were previously acquired from the Genetics of Hypertension Associated Treatments (GenHAT) study, an ancillary pharmacogenetics study of the Antihypertensive and Lipid-Lowering Treatment to Prevent Heart Attack Trial (ALLHAT).^27^ A total of 4526 GenHAT self-reported AAs were randomized to chlorthalidone, a thiazide diuretic, and had sufficient DNA for genotyping.

### Clinical data

BP and FG measurements in the ALLHAT study have been described.^28^ For the current study, BP response was defined as the change in systolic BP (SBP) and the change in diastolic BP (DBP) from baseline to 6 months post thiazide diuretic treatment initiation in the trial. FG response was defined as the change in FG from baseline to two years of a treatment since initiation. Other clinical data including body mass index and T2D status were measured at baseline. FG was not a primary outcome measured in ALLHAT and was missing for 71.9% of the sample at the 2-year visit. In total, 317 were missing data on change in DBP, 320 were missing data on change in SBP, and 3156 were missing data on change in FG.

### Genotyping and Quality Control (QC)

GenHAT samples were genotyped using the Expanded Illumina Infinium Multi-Ethnic AMR/AFR Beadchip, using the manufacturer’s protocol (Illumina, Inc., San Diego, CA). Genotypes were called using the Illumina GenomeStudio software. In total, 4296 samples were successfully genotyped, including 1559 mtDNA single nucleotide polymorphisms (SNPs). We set both the missing rate for samples and SNPs to 5% (removing 17 samples and 13 SNPs), and we removed 944 SNP duplicates, 261 singletons, 4 SNPs with potential heteroplasmic issues, and 26 SNPs with no dbSNP formally assigned rsID. We retained 4279 samples and 311 mtDNA SNPs to conduct a single SNP and haplogroup analysis. After these data were merged with the complete clinical data described above, there were 3962 (317 missing), 3959 (320 missing), and 1123 (3156 missing) samples for the analysis of DBP response, SBP response, and FG response with genotypes, respectively.

### Haplogroup Discovery

Haplogroups were obtained for each individual using HaploGrep,^29, 30^ an open-source maternal haplogroup classifier. According to the recommendation from HaploGrep’s developer, we removed a total of 704 samples due to call discrepancy with the classified haplogroup HV, H2, and Z7. After merging this data with the complete clinical data described above, there were 3310, 3308, and 959 samples for the analysis of DBP response, SBP response, and FG response with haplogroups, respectively.

### Statistical Methods

We standardized change in SBP and change in DBP outcomes by Z-transformation. FG was transformed using an inverse rank-based transformation. We used linear regression models to examine the association between individual mtDNA haplogroup clades L, M, and N and SBP, DBP, and FG response. Next, we examined the association of haplogroup L subclades with each response (DBP *n* = 2803; SBP *n* = 2801; FG *n* = 786) (L subclades sample count is a subset of the original Haplogroup sample set, that’s stated above (n= 3310)). The L3/L4 groups were combined due to small sample sizes (Figure 1). All models were adjusted for baseline measures (DBP, SBP, or FG), age, and sex. Using the Bonferroni correction, we calculated the level of significance for each outcome (DBP, SNP, and FG response against clades and subclades) to 0.05/ (3 outcomes) = 0.0167. Finally, we conducted a linear regression analysis to examine the association between individual mtDNA variants and SBP, DBP, and FG response under an additive genetic model adjusting for baseline measure (DBP, SBP, or FG), age, sex, and principal components to control for global ancestry (estimated using EIGENSTRATv6.1.4).^31^ Using the Bonferroni correction, we set the nominal significance to 0.000053 (=0.05/ (311 SNPs*3 outcomes)).

**Figure 1.**
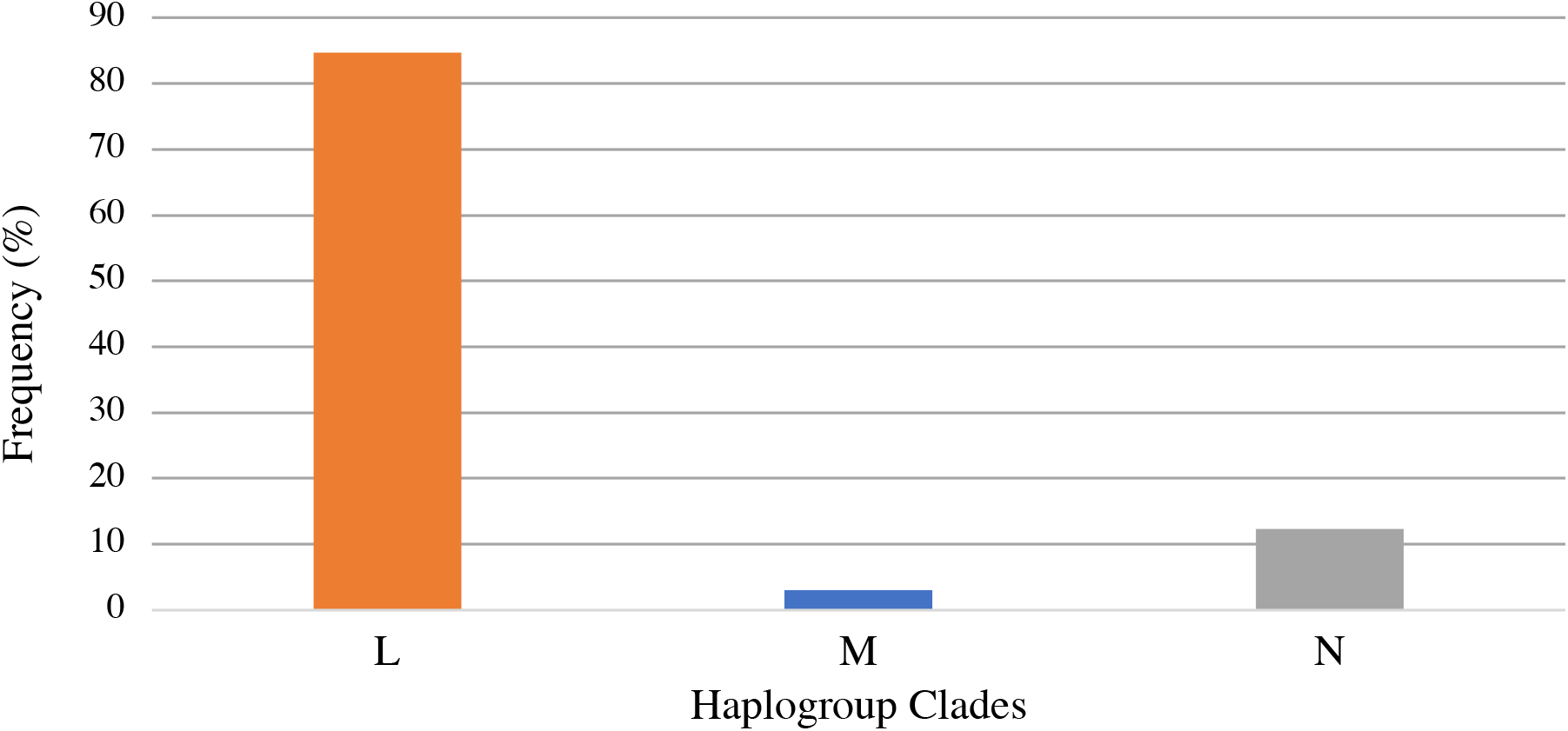
Sample Frequency Distribution of mtDNA Haplogroup Clades L, M, and N

## Results

### Clinical Data

Figure 1 shows the distribution of the mtDNA haplogroup clades L, M, and N in the GenHAT sample with clade L being the most common (84.68%). Table 1 presents the cohort baseline characteristics for the haplogroup analysis. The participants had a history of cigarette smoking (27.63%), ~41% had T2D, and ~14% were diagnosed with stage 3 or greater chronic kidney disease (Table 1). Mean (± standard deviation, SD) baseline SBP was 145.6 mmHg (±15.71 mmHg); DBP was 84.48 mmHg (±10.03 mmHg), and FG was 127.6 mg/dL (±65.93 mg/dL). Mean (±SD) SBP response was −6.51mmHg (±19.13mmHg); DBP response was - 3.01mmHg (±10.78 mmHg), and FG response was 6.95 mg/dL (±56.83 mg/dL) (Table 1).

**Table 1.** Baseline characteristics of GenHAT sample subset for Haplogroup Analysis

| Characteristics | Cases |
| --- | --- |
| Age, years | 66.1±7.73 |
| Female sex, n (%) | 1836 (55.47) |
| BMI, kg/m<sup>2</sup> | 30.4±6.75 |
| T2D, n (%) | 1351 (40.82) |
| CKD<sup>a</sup>, n (%) | 459 (14.40) |
| Cigarette smoker<sup>a</sup>, yes (%) | 772 (27.63) |
| Baseline SBP, mm Hg | 145.6±15.71 |
| Baseline DBP, mm Hg | 84.48±10.03 |
| Baseline FG, mg/dL | 127.4±65.93 |
| Baseline serum potassium (K<sup>+</sup>), mmol/L | 4.22±0.54 |
| SBP response, mm Hg | -6.51±19.13 |
| DBP response, mm Hg | -3.01±10.78 |
| FG response, mg/dL | 6.95±56.83 |
| BP: time between baseline and follow-up, months | 6 |
| FG: time between baseline and follow-up, months | 24 |

BMI, body mass index; T2D, type 2 diabetes; CKD, chronic kidney disease; SBP, systolic blood pressure; DBP, diastolic blood pressure; FG, fasting glucose; BP, meaning both SBP and DBP
a CKD was calculated as baseline eGFR $\leq 60$ mL/min/1.73m<sup>2</sup>
Continuous variables are shown as mean ± standard deviation and dichotomous traits are shown as *n* (%)

### Haplogroup Association Analysis

There were no significant associations between haplogroup clades (L, M, N) and change in SBP, DBP, or FG (Table 2). When examining the L subclades, we found no significant association of subclades with change in SBP and DBP. However, the 3 subclades (L1, L2, and L3/L4) compared to reference subclade L0 were significantly associated with change in FG (β= −0.405; −0.354; −0.365, P-value < 0.017), respectively (Table 2).

**Table 2.** Regression Analysis Results of Haplogroup Clades and subclades

| | $\Delta$ FG<sup>a</sup> | | $\Delta$ DBP<sup>b</sup> | | $\Delta$ SBP<sup>b</sup> | |
| --- | --- | --- | --- | --- | --- | --- |
| | $\beta$ (S.E.) | P-value | $\beta$ (S.E.) | P-value | $\beta$ (S.E.) | P-value |
| **Clades M and N vs. reference Clade L (DBP n = 3310; SBP n = 3308; FG n = 959)** |  |  |  |  |  |  |
| M | 0.226<br>(0.163) | 0.166 | -0.059<br>(0.083) | 0.480 | -0.110<br>(0.087) | 0.207 |
| N | 0.020<br>(0.088) | 0.822 | -0.072<br>(0.043) | 0.092 | -0.019<br>(0.045) | 0.669 |
| **Subclades L1, L2, L3/L4 vs. L0 (DBP n = 2803; SBP n = 2801; FG n = 786)** |  |  |  |  |  |  |
| L1 | -0.405<br>(0.134) | 0.003 | -0.004<br>(0.064) | 0.951 | 0.071<br>(0.669) | 0.286 |
| L2 | -0.354<br>(0.133) | 0.008 | 0.047 (0.062) | 0.448 | 0.107<br>(0.653) | 0.102 |
| L3/L4 | -0.365<br>(0.135) | 0.007 | 0.015 (0.064) | 0.816 | 0.064<br>(0.671) | 0.342 |

a Outcome variable $\Delta$ FG was rank-based inverse transformed
b Outcome variables $\Delta$ DBP and $\Delta$ SBP were standardized
All models were adjusted for age, sex, and baseline measure that matched the outcome (baseline FG, baseline DBP, and baseline SBP)

### Single Variant Association Analysis

No mtDNA SNP was associated with SBP, DBP, or FG response after correction for multiple testing. SNPs with marginal significance are presented in Table 3. In total, we observed 1 mtDNA SNP that was marginally associated with a change in SBP, 6 SNPs with a change in DBP, and 11 SNPs with a change in FG with P-value < 0.05.

**Table 3.** Association analyses of mtDNA SNPs with baseline DBP and FG

| ID | Trait | dbSNP ID | BP | MAF | BETA | SE | P | Genes* |
| --- | --- | --- | --- | --- | --- | --- | --- | --- |
| exm2216342-BOT-1 | SBP | rs1116906 | 8460 | 0.015 | -0.109 | 0.056 | 0.05204 | *ATP8* |
| Seq-rs28358578-BOT-2 | DBP | rs28358578 | 2332 | 0.111 | 0.061 | 0.021 | 0.0028 | *ND1* |
| exm2216198 | DBP | rs28358570 | 921 | 0.059 | -0.076 | 0.027 | 0.0054 | - |
| Seq-rs28359170-BOT-1 | DBP | rs28359170 | 12236 | 0.101 | 0.047 | 0.021 | 0.0297 | *ND4 - ND5* |
| Seq-rs28358573-BOT-2 | DBP | rs28358573 | 1442 | 0.101 | 0.0465 | 0.021 | 0.0300 | *ND1* |
| Seq-rs41352249-BOT-2 | DBP | rs41352249 | 6680 | 0.041 | -0.069 | 0.032 | 0.0314 | *COI* |
| Seq-rs202123618-TOP-1 | DBP | rs202123618 | 3434 | 0.018 | 0.101 | 0.048 | 0.0339 | *ND1* |
| Seq-rs2853502-BOT-2 | FG | rs2853502 | 13276 | 0.051 | 0.171 | 0.063 | 0.0096 | *ND5* |
| Seq-rs2853487-TOP-3 | FG | rs2853487 | 10589 | 0.054 | 0.169 | 0.062 | 0.0106 | *ND4L* |
| rs373606184 | FG | rs3020601 | 5442 | 0.055 | 0.153 | 0.061 | 0.0145 | *ND2* |
| Seq-rs200931747-BOT-2 | FG | rs193303033 | 4312 | 0.049 | 0.168 | 0.064 | 0.0151 | *ND1 - ND2* |
| Seq-rs2853489-TOP-1 | FG | rs2853489 | 1122 | 0.012 | 0.340 | 0.140 | 0.0158 | *ND4* |
| exm2216342-BOT-1 | FG | rs1116906 | 8460 | 0.015 | 0.304 | 0.121 | 0.0213 | *ATP8* |
| Seq-rs28357376-BOT-2 | FG | rs28357376 | 15824 | 0.068 | -0.125 | 0.053 | 0.0336 | *CYB* |
| Seq-rs28445203-BOT-2 | FG | rs28445203 | 1123 | 0.016 | 0.246 | .118 | 0.0372 | *ND1 - ND2* |
| exm2216469-BOT-1 | FG | rs35070048 | 1109 | 0.059 | -0.116 | 0.057 | 0.0428 | *CYB* |
| exm2216384 | FG | rs28358274 | 10086 | 0.068 | -0.116 | 0.053 | 0.0460 | *ND3* |
| Seq-rs2854125-BOT-2 | FG | rs201893071 | 1095 | 0.068 | 0.113 | 0.057 | 0.0479 | *CYB* |

\*Genes were annotated from Database of Single Nucleotide Polymorphisms (dbSNP [Sherry ST, et al. dbSNP: the NCBI database of genetic variation. Nucleic Acids Res, 2001; 29:308-311.])
(Gene – Gene: SNP is located between the 2 Genes.)
DBP, diastolic blood pressure; FG, fasting glucose; SNP, single nucleotide polymorphism; BP, base pair; MAF, minor allele frequency; BETA, parameter estimate from regression model; SE, standard error of BETA; P, SNP p-value

## Discussion

This study examined the association between mtDNA genetic variants and haplogroups and BP and FG response to a thiazide diuretic within a large sample of hypertensive AAs. Results from the single variant analysis did not remain statistically significant after correcting for multiple testing, however the haplogroup analysis suggested FG response may differ within the L subclade. The discoveries obtained here complement previous research showing association of mtDNA variation with cardiometabolic traits and importantly builds on pharmacogenetics research focused on first-line AHT response in AAs.^12, 32^

Consistent with other studies of this population, the majority of GenHAT samples were classified into the mtDNA haplogroup L clade (over 80% of the sample, see Figure 1). ^33,34^ When considering the major clades (L, M, N) in the GenHAT population, we did not observe an association with any of the response outcomes. However, within the L clade, subclades L1, L2, and L3/L4 (vs. L0) had an inverse association with change in FG suggesting a potential protective effect. While few studies have reported on the association of mtDNA haplogroups with FG changes during AHT treatment, some research suggests haplogroups are associated with T2D.^32, 35^ For instance, Sun and colleagues reported women living with HIV possessing haplogroup L2 (versus non-L2 groups of L0/L3/other) had a 49% lower risk of developing T2D, suggesting haplogroup L2 may potentially be a protective factor.^34^ Other reports of haplogroup association with T2D are mostly in other race groups. The haplogroups J/T and T were found to be associated with an increased risk of diabetes in Europeans.^36^ The M8a^37^, B4, and D4^38^ haplogroups have an association with T2D in East Asians. However, the haplogroup N9a^39^ has been suggested to confer resistance against T2D in the Japanese and Korean populations and to be a protective factor against metabolic syndrome in Japanese women.^40^ With multiple studies suggesting haplogroups may be related to T2D within race groups, more research on L subclades and FG levels during AHT treatment is warranted.

In the single variant analysis, we observed SNPs with marginal association to SBP, DBP, and FG response to chlorthalidone. These SNPs were in or near NADH-Ubiquinone Oxidoreductase Chain 1 (*ND1*), NADH-Ubiquinone Oxidoreductase Chain 2 (*ND2*), NADH-Ubiquinone Oxidoreductase Chain 4 (*ND4*), NADH-Ubiquinone Oxidoreductase Chain 5 (*ND5*), and Cytochrome B (*CYB*). Though few studies report on mtDNA SNPs and response to AHT treatment, researchers continue to discover significant associations between mtDNA variants and cardiometabolic traits across different ancestries (*e.g*. European^12, 13, 41^, and African^12, 41^). For example, Liu and colleagues reported a strong association between a nonsynonymous mtDNA variant (m.5913G>A in the cytochrome c oxidase subunit 1 of respiratory complex IV) and SBP and FG in a sample of 7000 European Americans from the Framingham Heart Study (minor allele being associated with a > 5 mm Hg and >15 mg/dL higher SBP and FG level on average, respectively).^13^ Similarly Buford and colleagues examined the association between mtDNA variants measured by sequencing and SBP and mean arterial pressure across several ancestries (European (*n* = 1739), African (*n* = 683), and others (*n* = 121)).^41^ They detected a statistically significant association of mtDNA variants with higher SBP (e.g. m.93 A>G in hypervariable segment II (HVII); P-value = 0.0001) and mean arterial pressure (e.g. m.16172T>C in a hypervariable segment I (HVI); P-value = 0.0002), respectively within African-ancestry samples.^41^ Interestingly, these specific variants were low frequency in the other race groups and not associated with these traits. Overall, these studies are supportive of a role for mtDNA variants in BP and FG; however, similar associations were not extrapolated to response to AHT treatment in our study.

This study is not without limitations. First, the coverage of mtDNA variants was restricted to GWAS data. Overall, there are no “gold standard” methods for calling mtDNA haplogroup with GWAS data and we used standard methodology adopted by another study for the same GWAS array.^42^ Ultimately, deeper genomic coverage may enhance SNP and haplogroup detection using available software. Further, FG was missing on a considerable number of samples in our study. This is a limitation of the ALLHAT study since BP and cardiovascular disease outcomes were the main focus. Still, several published studies have made use of the available glucose data in this large, NIH-funded clinical trial.^5, 43–45^ Despite these limitations, this study remains the largest AA pharmacogenomics study to examine mtDNA variants and haplogroups with BP and FG response.

In summary, this study represents the most extensive investigation of the role of the mitochondrial genome in BP and FG response to chlorthalidone (a thiazide diuretic) within AAs. The overwhelming percentage of AAs suffering from HTN reveals the urgency to discover causal determinants of better BP response to treatment. Despite the urgency, there remains a lack of pharmacogenomic studies within hypertensive AAs. Hence, this study provides the inclusive research that is required to aid understanding of AHT treatment response and eventually facilitate better blood pressure control. The discoveries presented in this study support future investigations of mitochondrial genomics in metabolic response to chlorthalidone among AAs.

## Acknowledgements

The authors are thankful to both ALLHAT and GenHAT for the use of their datasets. The genotyping was funded by R01HL123782.

## Discloser Statement

The authors disclose no conflict of interest.

## Data Availability Statement (DAS)

The data that support the findings of this study are available from the corresponding author, BM, upon reasonable request.

